# Similar somatotopy for active and passive digit representation in primary somatosensory cortex

**DOI:** 10.1101/754648

**Authors:** Zeena-Britt Sanders, Daan B Wesselink, Harriet Dempsey-Jones, Tamar R Makin

**Affiliations:** Wellcome Centre of Integrative Neuroimaging, FMRIB, John Radcliffe Hospital, Headington, Oxford OX3 9DU, United Kingdom; Institute of Cognitive Neuroscience, University College London, 17 Queen Square, London WC1N 3AZ

**Keywords:** Topography, somatosensation, finger representations, hand map, motor

## Abstract

Scientists traditionally use passive stimulation to examine organisational properties of the primary somatosensory cortex (SI). Recent research has, however, emphasised the close and bidirectional relationship between somatosensory and motor systems. This suggests active contributions (e.g., direct inputs from the motor system to SI) should also be considered when studying SI representations. Under such a framework, discrepant results are possible when different tasks are used to study the same underlying somatosensory representation. Here we examined whether SI hand representation is consistent when compared between active and passive tasks. Moreover, we ask whether this holds when task demands and stimulus properties are not directly matched. Using 7T fMRI, we examined three key digit representation features – spatial location, activity gradient profiles and multivariate representational structure. We show that although activity was increased overall in the active task, all key features of digit somatotopy were consistent over tasks, using both traditional univariate (activity-level) and multivariate (pattern structure) measurements. Despite overall comparability, when investigating fine-grained features of somatotopy using multivariate analysis, the active task was superior for individuating the representation across digits. We suggest that the information produced by an active task is more aligned with typical, ecologically relevant patterns of hand sensory input, resulting in more optimal SI activity patterns. Our findings validate the utilisation of active tasks for studying SI somatotopy.

## 1. Introduction

Traditionally, the organisation of the somatosensory system has been studied using passive tactile stimulation protocols, e.g., contacting stationary digit(s) with tactile stimuli. While providing a high controlled means to study the primary somatosensory cortex (SI; Besle, Sánchez-Panchuelo, Bowtell, Francis, & Schluppeck, 2014; Merzenich, Kaas, Sur, & Lin, 1978; Sanchez-Panchuelo, Francis, Bowtell, & Schluppeck, 2010; Sur, Merzenich, & Kaas, 1980), such protocols may provide a constrained view of somatosensory inputs as they habitually occur. For instance, there is growing recognition that somatosensory and motor processing are intimately and bi-directionally linked (for example: Darainy, Vahdat, & Ostry, 2013; Pruszynski et al., 2016; Scott, 2004). In daily life, the majority of hand input to somatosensory cortex occurs as a result of deliberate action (i.e., is driven by motor control), or otherwise is directly relevant for supporting motor control (Scott, 2004). Accordingly, recent research shows that the organisation of somatosensory hand representation reflects habitual patterns of cooperative action between digits (Ejaz, Hamada, & Diedrichsen, 2015; Ingram, Körding, Howard, & Wolpert, 2008). These usage effects are manifested in greater representational overlap between digit representations (e.g., multi-digit receptive fields) for digits used more frequently together in daily life (Ejaz et al., 2015; Merzenich et al., 1984). These findings suggest that motor synergies dominating everyday hand use are reflected in SI digit representation.

As a further rationale that passive tactile stimulation may provide a restricted view of somatosensory activity, the somatosensory cortex receives inputs from a host of sources beyond the simple cutaneous touch activated by passive tasks. This includes activation of deep mechanoreceptors, muscle proprioceptors, and skin stretch receptors (Bensmaia & Tillery, 2014; Chouvardas, Miliou, & Hatalis, 2008; Saal, Delhaye, Rayhaun, & Bensmaia, 2017). Active movement also causes efferent signals into SI from the motor system (Wolpert & Flanagan, 2001; Wolpert, Ghahramani, & Jordan, 1995), as well as top down inputs from high-order cognitive processing, e.g., attention, visual input (Kuehn, Mueller, Turner, & Schütz-Bosbach, 2014; Puckett, Bollmann, Barth, & Cunnington, 2017). In our daily life, this multitude of inputs are meaningfully integrated in our actions and interactions with the world. Therefore, there is a growing interest to study the somatosensory system under active conditions, which provide a richer and arguably more ecologically-relevant model for somatosensory processing.

However, biomechanical constraints during movement could drive differences in SI activity profile during active versus passive tasks. Body parts range in their capacity for independent movement (known as ‘enslavement’; Lang & Schieber, 2004; Reilly & Schieber, 2003). Higher inter-dependence in movement could limit the possibility for active tasks to effectively delineate the representations of enslaving/enslaved body parts, thus preventing fine-grained maps. As an example, in a single digit tapping task, movement of digit four will activate skin stretch and deep proprioceptors in the palm, as well as causing some degree of enslaved movements in digits three and five (and the resulting sensory inputs from these movements). Aligning with this idea, active tasks have been previously demonstrated to produce greater overall SI activation compared to passive stimulation (Berlot, Prichard, Reilly, Ejaz, & Diedrichsen, 2019; Wiestler, Mcgonigle, & Diedrichsen, 2011). Whilst these concerns are valid, it may also be the case that these biomechanical constraints influence the basic SI organisation, and that through incorporating active tasks to study SI processing we can shed light on these organisational principles (as elaborated in the Discussion below). Despite these conceptual and empirical differences, few studies have directly compared whether key features of somatotopy (such as digit selectivity and overlap) remain consistent between active and passive tasks.

The hand representation in SI provides an ideal test case for the stability of somatotopic mapping under active and passive tasks. This area contains detailed digit maps, with physically adjacent digits represented next to each other. Using high-field neuroimaging, it is now possible to identify these characteristic digit maps in humans, with high inter- and intra-participant reliability (Ejaz et al., 2015; Kolasinski et al., 2016). Digit somatotopy is characterised in neuroimaging by two main principles: 1) digit selectivity, meaning separable regions show increased responsiveness to one digit compared to all other digits (Kolasinski et al., 2016), and 2) inter-digit overlap, where neighbouring digits are more overlapping in their representation than non-neighbouring digits (Ejaz et al., 2015). The features of these digits maps have been extensively studied separately using either active (Kikkert et al., 2016; Wesselink et al., 2019) or passive tasks (Martuzzi, van der Zwaag, Farthouat, Gruetter, & Blanke, 2014; Sanchez-Panchuelo et al., 2010; Schweizer, Voit, & Frahm, 2008). Recent work by Berlot and colleagues (2019) provided a comparison between active and passive tasks. This study has shown that when task and stimulus properties are tightly matched, active and passive stimulation produce digit specific representations to the same extent, despite higher activity levels in the active condition. Specifically, the average dissimilarity between the digits’ activity patterns in contralateral SI, a multivariate measure indicating inter-digit decodability (see Methods), was not significantly different between conditions. Still, it remains to be determined whether these results are achieved due to the matched sensory and motor demands, or whether representation will diverge given very different task demands. Moreover, the multivariate dissimilarity measure above is naïve to spatial relationships. It is, thus, unable to inform on the critical properties of spatial correspondence and the somatotopic gradient of activity levels, which has been the focus of most previous studies (e.g. Flor et al., 1995; Makin, Scholz, Henderson Slater, Johansen-Berg, & Tracey, 2015). For instance, multivariate measures reveal similar hand representational structure across SI and MI (Berlot et al., 2019), though SI but not M1 is known to have highly-ordered digit somatotopy (Huber et al., 2018).

Here we use 7T fMRI to investigate whether critical features of SI hand somatotopy – spatial location, inter-digit activity profiles and multivariate representational structure (defined in detail in the Methods), are preserved for a task providing passive cutaneous stimulation, and a haptic motor task. While these tasks afford different forms of sensory feedback, hand position, motor signals, task demands, and attentional requirements, we hypothesised that resulting digit somatotopy would be similar, because despite differing types or amount of input, all inputs feed into somatotopically restricted regions (Kuehn et al., 2017; Qi & Kaas, 2004; also see Discussion). However, consistent with Berlot et al., (2019), increased somatosensory input in the active condition was predicted to lead to greater overall activity in the active task, compared to the passive.

## 2. Methods

### 2.1 Participants

Fifteen healthy volunteers (6 female, age ± S.E.M. = 26.44 ± 1.04) were recruited for this study. All participants except one were right handed and all experimental tasks were performed using the right hand. The fMRI session was part of a larger study, full information and study protocol can be found in https://osf.io/nh4yp/. All participants gave written informed consent and ethical approval for the study was obtained from the Health Research Authority UK (13/SC/0502). One participant was excluded as an outlier from the passive multivariate task analysis (>3σ below group mean). This participant was included in the active condition where their results were within normal group variance (0.3σ above group mean).

### 2.2 MRI Tasks

#### 2.2.1 General procedure

Participants completed an active and a passive task whilst in the scanner during a single session (order counter-balanced between participants). These tasks were completed in separate scans (four of each), with short rest breaks in-between. The general structure of the scans was identical for the passive and active tasks. Each scan comprised of a pseudo-randomised block design, with each of the five digit stimulations repeated over three 12 second blocks as separate conditions, and 3 rest blocks of 12-24 sec. Four scans were carried out of each task. A functional localiser to independently identify digit-selective regions of interest was also conducted before the active task. This localiser was also organised into digit-specific blocks over two scans, but with a set inter-digit sequence design (travelling wave, also known as phase-encoding design; Kikkert et al., 2016; Kolasinski et al., 2016; Mancini, Haggard, Iannetti, Longo, & Sereno, 2012; Sanchez-Panchuelo et al., 2010; Wandell, Dumoulin, & Brewer, 2007; Zeharia, Hertz, Flash, & Amedi, 2015).

#### 2.2.2 Active task

During the active task, participants were required to move individual digits by making button presses on a MRI-safe keyboard with five keys (Berlot et al., 2019). The digit to be moved in the upcoming block was indicated by a visual display of five grey bars, with one highlighted green – indicating the digit to be moved during the upcoming block (see figure 1 left). In each trial participants were required to move a cursor from the starting position (inside a grey box at the lower part of the screen) into a green box that appeared above the grey box by pressing the relevant keyboard key. After hovering the cursor inside the box for 400ms they received a point for successful completion, and were instructed to release the force on the keyboard key – allowing the cursor to fall back into the starting position grey box, ready for the next trial. Participants completed 12 trials per digit before receiving instructions for the next digit. The keys required force application to activate, though the actual displacement of the digit was low (∼10mm).

**Figure 1:**
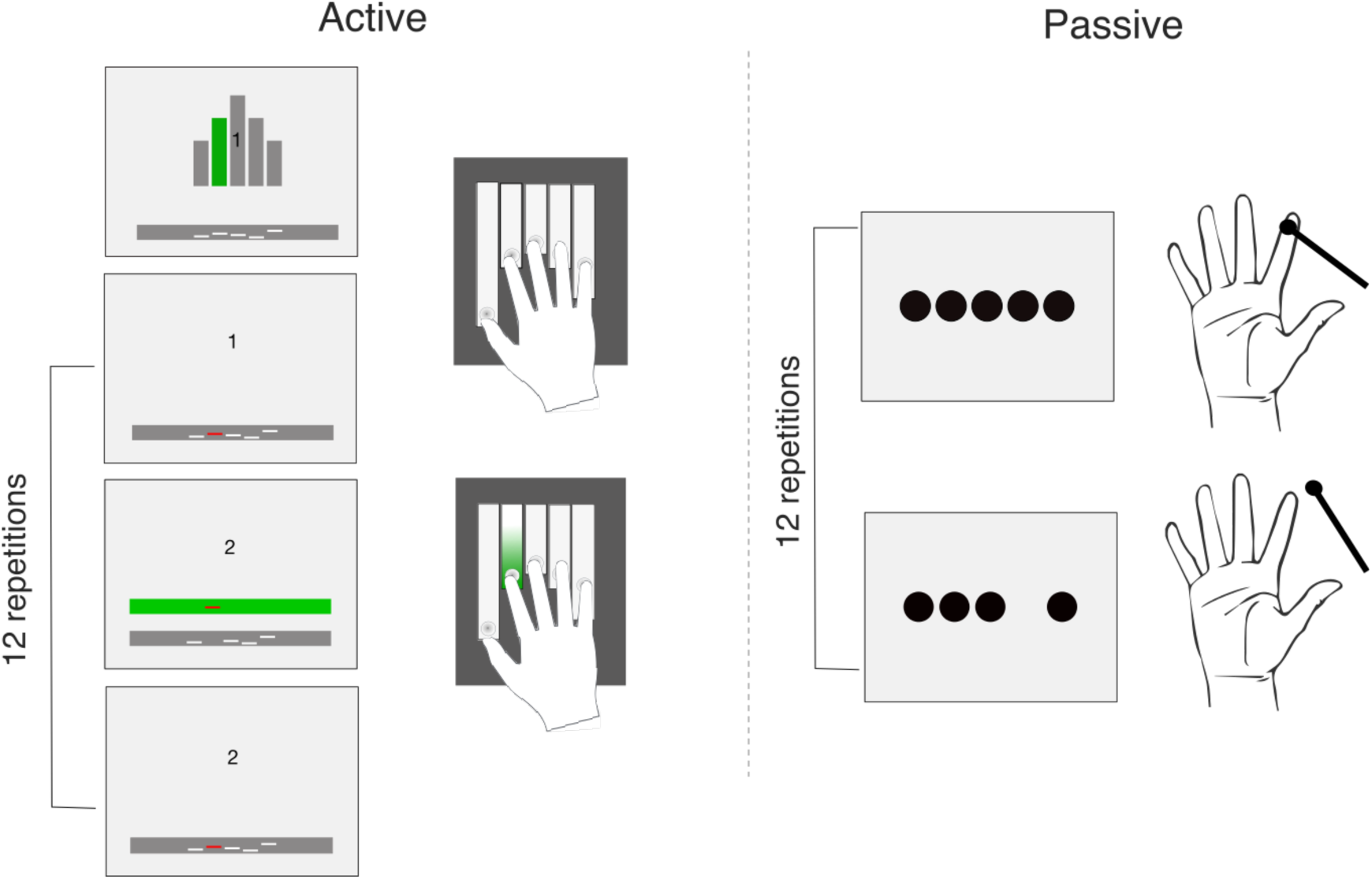
Overview of the two tasks used in this study. During the active task (left) participants were required to press buttons on a MRI-safe keyboard with their right hand. Five white lines within a grey box near the bottom of the screen represented the five digits at rest. The digit to be used in the upcoming trial was highlighted in green on a hand schematic (top). The line of the digit to be used was then coloured red. Participants were instructed that whenever the green box appeared above the grey, they were to press the key and move the red line into the green box for 400ms before releasing. Participants were awarded points for successful completing each trial. In the passive task (right), participants rested their right hand in a supine position on a foam support. Participants had their individual digits probed by the experimenter at a rate of once per second. At the same time, they saw five dots displayed on the screen corresponding to their five digits and the dot corresponding to the digit being stimulated would flash on and off at the same time as the stimulation was occurring.

#### 2.2.3 Passive task

In the passive stimulation task, participants were asked to rest their right hand in a comfortable, supine position on a foam support. The experimenter used a plastic probe to stimulate each digit by tapping the probe against the distal pad of the digit at a 1Hz rate for twelve seconds, before moving on to the next digit. During the task participants were shown five white dots on the screen, corresponding to each of the five digits (figure 1 right). The dots flashed individually at a rate of 1/second whilst the digit was being stimulated. To promote engagement across the duration of the task, double taps would be administered randomly (one double-tap per digit condition in each session). Participants were asked to indicate when they felt the double taps by pressing a button on a button-box placed in their left hand.

#### 2.2.4 Digit-selective clusters localiser

A travelling wave task was used as an independent functional localizer for digit-specific regions-of-interest (ROIs). The travelling wave protocol involves a set digit conditions cycle. This approach is particularly useful to selectively locate voxels that show an enhanced response to one digit compared to all other digits, and has previously been used to identify SI digit somatotopy (for example: Kikkert et al., 2016; Kolasinski et al., 2016; Sanchez-Panchuelo et al., 2010).

Participants used the same keyboard and visual display as in the active task (see section 2.2.2). Two separate scans with reverse orders (forward and backward cycles) were used to overcome potential order-related biases due to the sluggish hemodynamic response. In the forward scan, the order of digit blocks cycled from digit 1 to digit 5 (D1-D2-D3-D4-D5) whereas the reverse order was used in the backward scan (D5-D4-D3-D2-D1). In each scan, the cycle was repeated five times continuously with no rest periods in between. During the cycles, each digit was moved 8 times (at 1Hz) before the instructions for the next digit were shown. As in the active task, at the start of each block, the digit to be used in the upcoming block was visually cued (see figure 1), then 8 digit presses were made, before receiving the cue for the next digit.

### 2.3 MRI acquisition

All MRI measurements were acquired using a Siemens 7 Tesla Magnetom scanner with a 32-channel head coil. Task fMRI data was acquired using a multiband EPI sequence with an acceleration factor of 2 (Moeller et al., 2010; Uǧurbil et al., 2013). A limited field-of-view was used consisting of 56 slices each 1mm thick over the primary somatosensory cortex with a 192×192mm in-plane FOV (TR 2000ms, TE 25ms, FA 85deg, GRAPPA 3). This resulted in spatial resolution of 1mm isotropic. A whole brain anatomical T1-weighted image was also collected with a 1mm isotropic spatial resolution (TR 2200ms, TE 2.82ms, FA 7deg, TI 1050ms).

### 2.4 MRI pre-processing

All MRI data pre-processing and analysis was carried out using FMRIB Software Library (FSL; version 6.0) as well as Matlab scripts (version R2014b) which were developed in-house. Surface reconstruction was carried out using Freesurfer (Dale, Fischl, & Sereno, 1999) www.freesurfer.net) and results from the task and travelling wave analysis were projected onto the cortical surface for visualization purposes using Connectome Workbench software (www.humanconnectome.org).

#### 2.4.1 Pre-processing

Standard pre-processing steps were carried out using FSL (Jenkinson, Beckmann, Behrens, Woolrich, & Smith, 2012). FSL’s Expert Analysis Tool (FEAT) was used to carry out motion correction (using MCFLIRT; Jenkinson, Bannister, Brady, & Smith, 2002) brain extraction (BET; (Smith, 2002) spatial smoothing using a 1mm full width at half maximum (FWHM) Gaussian kernel and high pass filtering using a cut-off of 100s. The output from the MCFLIRT analysis were visually inspected for excessive motion (defined as > 1mm absolute mean displacement). No participants had an absolute mean displacement greater than 1mm.

#### 2.4.2 Image registration

For each participant, a midspace was calculated between the four active and four passive scans, i.e. the average space in which the images are minimally reoriented. Each scan was then aligned to this session-midspace using FMRIB’s Linear Image Registration Tool linear registration (FLIRT; Jenkinson et al., 2002; Jenkinson & Smith, 2001).

This registration was also run separately for the functional localiser (travelling wave) scans, where the forward and the backward scans were realigned to their midspace. The localiser midspace was then registered to the midspace of the active and passive tasks with using FLIRT.

Finally, as these scans were collected as part of a larger project containing two scanning sessions (https://osf.io/nh4yp/), the midspaces of the two scanning sessions were aligned together into a study-midspace. This was achieved by registering each of the individual session-midspace’s to the midspace of the structural scans collected during each session. Manual adjustments were made when needed to ensure an accurate co-registration of the hand-knob. Once an accurate registration was achieved, a final midspace was calculated from the sixteen scans across the two sessions and all scans were aligned to this study-midspace.

### 2.5 Initial analysis

#### 2.5.1 Active and passive tasks

A voxel-based general linear model (GLM) analysis was carried out on each of the 8 scans (4 active and 4 passive) using FEAT, to identify activity patterns for each digit condition. The design was convolved with the double-gamma haemodynamic response function, as well as its temporal derivative. Eleven contrasts were set up: each digit versus rest, each digit versus all other digits and all digits versus rest. The estimates from the four active/passive scans were then averaged voxel-wise using a fixed effects model with a cluster forming z-threshold of 3.1 and family-wise error corrected cluster significance threshold of P < .05, creating eleven main activity patterns for each task. These contrasts were used for the key analyses described below (section 2.6).

#### 2.5.2 Localiser & Regions-of-interest definition

The travelling wave approach described in section 2.2.4 was used in order to identify digit-specific voxel clusters. The analysis was carried out as in Kikkert et al. (2016). In short, a reference model was first created using a convolved hemodynamic response function to account for the hemodynamic delay. This model consisted of an 8 second ‘on’ period followed by 32 second ‘off’ period to model the movement block of one digit for one cycle. The model was shifted 20 times by one lag of 2 seconds (runs were acquired with a TR of 2s) in order to model one entire movement cycle (which lasted 40s), thus resulting in 20 reference models, this was repeated five times to model the five cycles in each scan. Following this the pre-processed BOLD signal time course for each voxel was correlated with each of the reference models. This resulted in cross correlation r-values, which were standardized using the Fisher’s r-to-z transformation. Lags were assigned to each digit (four lags per digit) in order to average the r-values across scans for each voxel. This resulted in a r-value for each digit which was further averaged across the forward and backward runs. To define digit specificity, each voxel was assigned to one digit using a ‘winner-takes-all’ approach. This was done by finding the maximum correlation for each voxel across the five averaged values, and assigning the voxel to the corresponding digit.

To correct for multiple comparisons, a false discovery rate (Benjamini & Hochberg, 1995) threshold (q < 0.01) was applied to each digit individually (as in Kikkert et al., 2016; see supplementary materials for average cluster size, and results from the same analysis using a more conservative threshold). The resulting FDR corrected digit-specific voxels were then used to create digit-specific clusters showing strong digit selectivity for each of the digits. This was done by using an anatomically defined mask of the hand region (SI hand mask, Figure 2A inset) which was defined for each participant based on a Freesurfer structural segmentation of SI subdivisions. Broadmann Areas 3a, 3b and 1, spanning a 2cm strip medial/lateral to the anatomical hand knob was included the mask (Wesselink et al., 2019). The digit specific activity under this mask was used to create digit-specific clusters (see figure 3A for example). This approach allowed us to identify digit-specific clusters on an individual participant basis, which were used for further analysis of the active and passive tasks.

**Figure 2:**
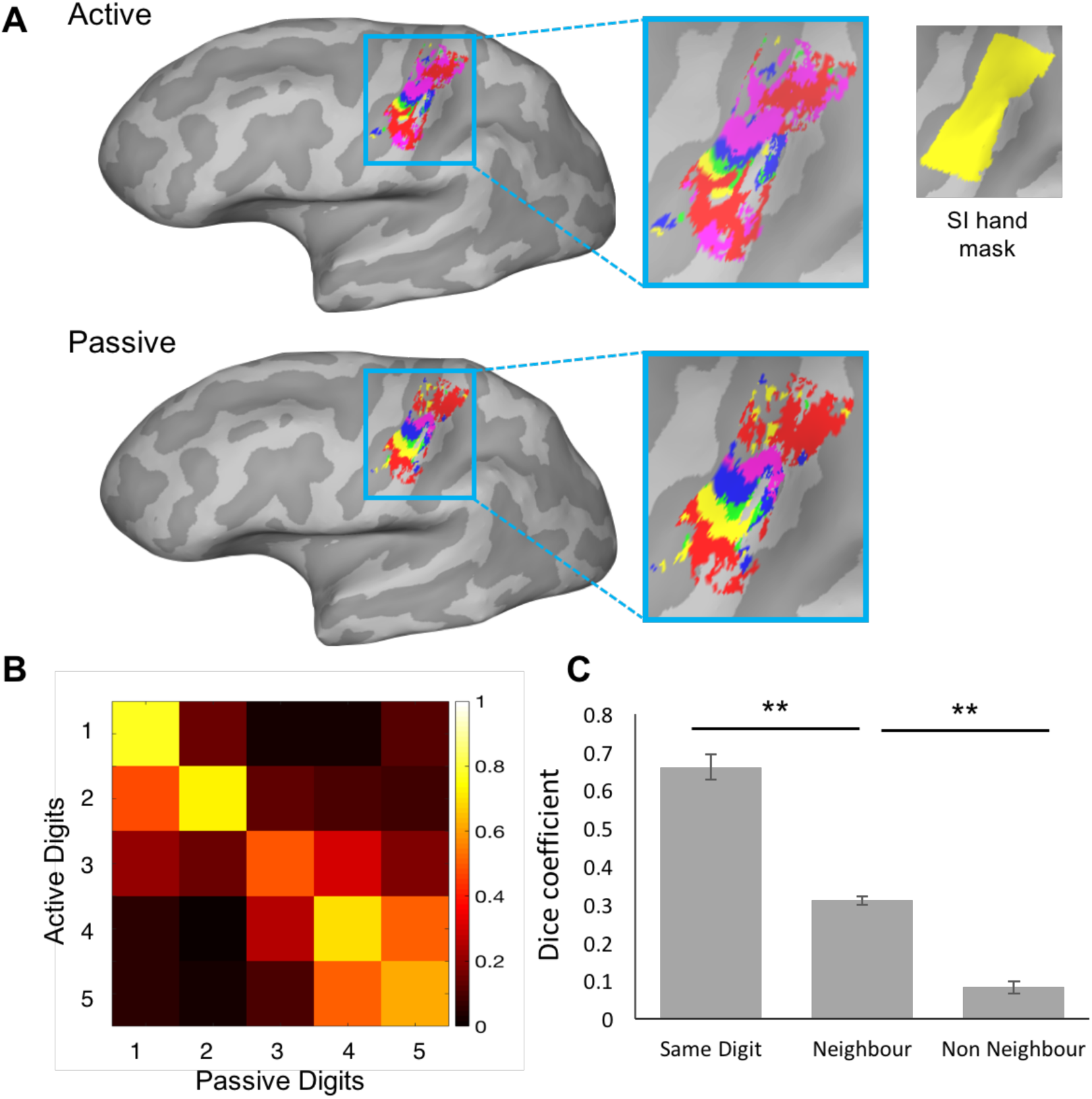
Overlap across active and passive tasks. (A) For an example participant, the minimally thresholded digit activity of each digit for both tasks is shown projected onto the cortical surface (red = D1, yellow = D2, Green = D3, Blue = D4, Pink = D5). Activity was masked using an anatomically defined SI hand mask including Broadmann areas 3a, 3b and 1, based on freesurfer segmentation, shown in inset. These activity maps were used for the Dice analysis. Note: Double D1 representation can be clearly seen in this participant. (B) 5×5 matrix showing spatial overlap in SI hand mask as measure by the Dice coefficient between active and passive digits. The diagonal represents the same digit overlap across tasks, whereas the offdiagonal elements represent overlap between neighbouring and non-neighbouring digits. Dice values are indicated in the bar, where hotter colour indicates greater overlap. (C) Bar chart showing average overlap for the same digits (averaged values across the diagonal line in (B)), neighbouring digits (averaged values of the offdiagonal in (B)), and non-neighbouring digits across the tasks (all other cells in (B)). The same digit across tasks has higher spatial overlap than neighbouring digits, which in turn showed higher overlap than non-neighbouring digits, as indicated in a ttest. Error bars show standard error of the mean. Significance is indicated by: ** = p<.001, * = p<.05.

**Figure 3:**
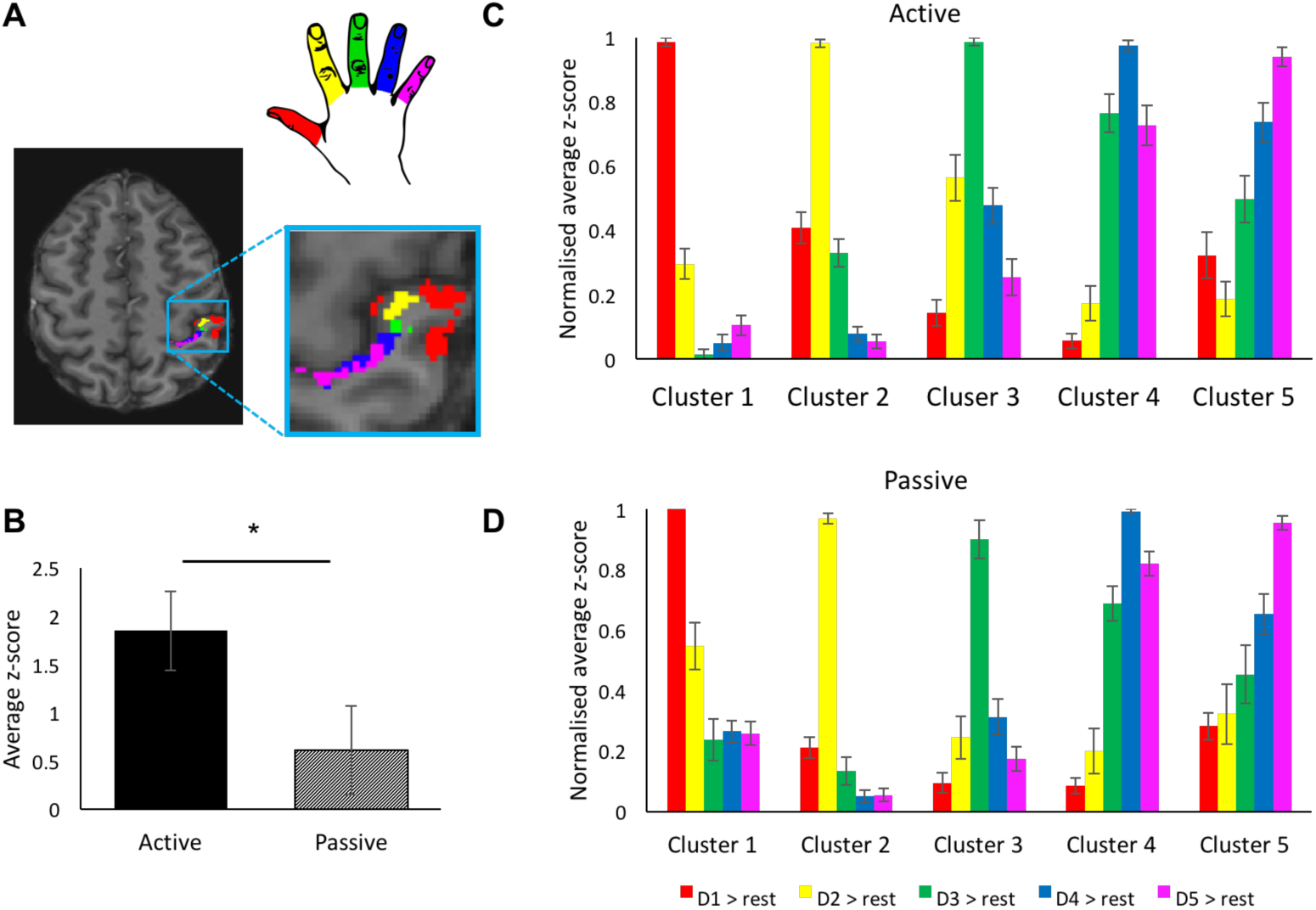
Somatotopic activity patterns across digit-selective clusters in active and passive tasks. A) Example of digit-specific clusters created using the travelling-wave approach, of one participant. Digit-specific clusters are predominantly located in Broadmann area 3b. Activity in each digit-specific cluster for each digit (versus rest) during the active or passive task was extracted. B) Activity was on average significantly higher in the active task than in the passive task. Therefore, for further analysis activity was normalised for each participant per digit. (C-D) Normalised activity (averaged across participants) for the (C) active and (D) passive tasks is shown in each digit-specific cluster (error bars show standard error of the mean). In both tasks, a somatotopic activity profile can be seen with each digit-selective cluster, with the target digit producing on average the greatest activity within its own digit-selective cluster, followed by its neighbouring digits, and lowest activity seen in non-neighbouring digits. Raw values (non-normalised) can be seen in figure S3. * indicates significant differences at p<.05.

### 2.6 Key analyses

All data used for final analysis will be made available online at the Open Science Framework upon publication of this manuscript.

#### 2.6.1 Spatial correspondence across tasks: Dice analysis

To assess the spatial correspondence between digit representations in the active and passive tasks, a Dice coefficient (Dice, 1945) was calculated on minimally-thresholded activity maps projected onto the cortical surface (as in Kikkert et al., 2016) for each individual participant. Traditionally, the amount of spatial overlap between, in this case, two digit representations, is calculated in reference to the total spatial area of these two digits. However, cases may occur when one digit’s representation is completely within another’s (therefore completely overlapping), but the area of the second digit may be much larger than the first, therefore leading to a small Dice coefficient. To account for these differences, the Dice coefficient was normalising to the smallest area of the pair of digits. This constrains the maximum overlap to be equal to the smallest digit representation area as shown below: (note - the same analysis was run with the traditional Dice coefficient with comparable results. This analysis can be found in the supplementary materials, figure S2).

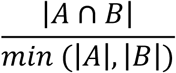

A represents the spatial area of one digit representation and B the spatial area of another. This produces values of spatial overlap ranging from 0 to 1, with 0 indicating no spatial overlap and 1 representing perfect overlap.

For each digit, the activity maps from the active and passive task (contrasting each digit vs all the other digits; see section 2.5.1) were projected onto the cortical surface, were minimally thresholded (Z>2) on the cortical surface and masked using the SI hand mask described above (see Figure 2A and Figure S1 for example participants). The spatial overlap between digit pairs was calculated across tasks. If the digit maps are overlapping, we should see more overlap between same vs. neighbouring digits, and between neighbouring than non-neighbouring digits. Paired-samples T-tests were carried out to determine whether this is the case across participants. Alpha levels were adjusted to account for these two comparisons (p<0.025).

#### 2.6.2 Somatotopic gradient in digit-specific clusters

We next looked at single-digit activity profiles in order to investigate whether the digit representations followed a comparable somatotopic relationship across tasks. To explore this, we looked at activity patterns for each digit within each digit specific cluster (identified with the independent localiser, see Figure 3A for an example) versus rest. For each digit the standardized activity vs. rest was extracted and then averaged for each digit specific cluster (Figure 3C-D). As overall activity varied across tasks (see figure 3B), normalised activity values were used for this analysis (see figure S3 for non-normalised values). Normalisation was carried out for each participant for both tasks and all digits, and was based on the highest activity value obtained for each digit across clusters.

To investigate the somatotopic gradient of activity within each digit-specific cluster, the difference in activity was calculated between the target digit (a single digit within its own digit-specific cluster) and its neighbours (e.g. in cluster 3: D3 vs D2 and D4) and non-neighbours (e.g. in cluster 3: D3 vs D1 and D5) across the two tasks. If activity follows a somatotopic gradient we would expect differences to be greater when comparing the target digit and its non-neighbours, than compared to its immediate neighbours. This allowed us to compare the somatotopic gradient in relation to the target digit, across active and passive tasks. We repeated the same calculation in each digit specific cluster for the non-target digits (e.g. in cluster 3: D4 vs D5 and D4 vs D1), allowing us to look at the somatotopic gradient for the non-target digits in each cluster (see figure S4 for all pairings).

Differences in somatotopic gradient (i.e. the differences between neighbouring or non-neighbouring digits) across the two tasks was compared for both the target and non-target digits using a repeated measures ANOVA. Significant interactions were followed up using post-hoc tests. As deviations from normality were revealed using the shapiro-wilks test for the residuals of two of the eight outcome variables, follow-up tests were instead conducted using non-parametric alternatives, in this case the Wilcoxon paired-ranks test. This was used to compare differences between neighbouring and non-neighbouring digits within tasks, as well as comparing neighbour and non-neighbours across tasks. Alpha levels were adjusted for these four comparisons (p<0.0125) The ANOVA has previously been found to be relatively robust to violations of normality (Glass, 1972; Harwell, 1992) and was therefore still carried out.

#### 2.6.3 Multivariate representational structure: Representational similarity analysis

We used representational similarity analysis (Kriegeskorte, Mur, & Bandettini, 2008) in order to assess the multivariate relationships between the activity patterns generated across digits and tasks. The (dis)similarity between activity patterns within the entire SI hand mask was measured for each digit pair using the cross-validated squared Mahalanobis distance (Nili et al., 2014). This was applied using the FSL-compatible toolbox in Wesselink & Maimon-Mor (2017). First, the activity patterns were pre-whitened using the residuals from the GLM and then cross-nobis distances were calculated for each task (active/passive) separately, using each pair of imaging runs and averaging those results. Due to cross-validations, the distance value is expected to be zero (but can go below) if two patterns are not statistically different from each other. Otherwise, greater distances indicate larger differences in multivariate representation.

The above analysis produced 10 distance values (one for each digit pair) per task, forming a representational dissimilarity matrix (RDM) per participant. Strength of representation was assessed by calculating the average value of the distances in an RDM (i.e., averaging across the values in the RDM, discounting values on the diagonal). Shape of representation was assessed by calculating the Spearman’s rho correlation between a measured RDM and the groups’ RDM (for each of the two tasks; mean excluding participant-of-interest; and RDMs were normalised prior to averaging not to bias the mean towards more dissimilar RDMs).

As an aid to visualise the RDMs (and not used in any statistical analysis), we also performed multidimensional scaling (MDS). This analysis projects the higher-dimensional RDM into a lower-dimensional space while preserving the inter-digit distances as accurately as possible (Borg & Groenen, 2003). MDS was performed on individual RDMs and averaged after Procrustes alignment (without scaling) to remove any arbitrary rotations introduced by MDS. Alignment was based on RDMs including both active and passive conditions (as well as a rest condition) as not to bias the visualisation. Dimensions were sorted on the basis of high variance between digits to demonstrate hand representation at the expense of non-digit-specific variation from rest.

## 3 Results

### 3.1 Spatial correspondence is observed across tasks

To quantify the extent of spatial overlap across the active and passive tasks, we calculated the Dice coefficient (Dice, 1945; see section 2.6.1) across digit maps. Minimally thresholded contrast maps for each digit vs baseline were compared within the SI hand mask encompassing BA 3a, 3b and 1 (see Figure 2A and Figure S1 for individual examples). Dice values were computed across tasks and digits resulting in a 5×5 matrix (figure 2B), with the diagonal representing the Dice coefficient of the same digits across active and passive tasks. We found that spatial overlap was maximal for comparison of the same digit between tasks (mean Dice = .660; see Figure 2C), compared with neighbouring (mean Dice = .311, t(14)=15.360 p< 0.001). Moreover, neighbouring digits showed greater overlap across tasks than non-neighbouring digits (mean Dice = .082, t(14)=10.948,p<.001). This analysis produces first evidence for spatial overlap in digit maps across tasks.

### 3.2 Activity profiles in both tasks show a consistent somatotopic gradient

We next wished to interrogate activity level profiles within and across digit-selective regions to investigate the somatotopic gradient of activity. For this purpose, we created digit-specific clusters using an independent localiser, designed to identify strong digit preference in SI (example: figure 3A). Please note: the nomenclature ‘target digit’ will be used to define activity related to a single digit within its own digit-specific cluster.

Average activity levels underlying each of the 5 clusters for each of the five digits was derived for each of the tasks (Figure S3 for raw values).

First, we wanted to confirm digit selectivity across paradigms and tasks was consistent. We found that in general there was a strong correspondence for digit-selectivity in the two tasks, with respect to the localiser cluster identity: The target digit showed the highest activity within its digit-specific cluster in 92-93% of the cases (69/75 “hits” for active and 70/75 for passive; Table 1).

**Table 1.** Proportion of participants showing winner-takes-all digit selectivity in accordance with the digit-specific clusters from the travelling wave paradigm.

| **Travelling wave** | **Active** | **Passive** |
| --- | --- | --- |
| D1 | 15/15 | 15/15 |
| D2 | 13/15 | 15/15 |
| D3 | 13/15 | 14/15 |
| D4 | 14/15 | 14/15 |
| D5 | 14/15 | 12/15 |

We next wished to compare the somatotopic cascade in activity across the two tasks. As clearly demonstrated in Supplementary Figure 3, activity levels were greater in the active task in comparison to the passive task, as confirmed in a paired t-test (averaged activity across all ROIs in the two tasks; t(14)=2.85, p=0.013). This result could be confounded by the fact that the travelling-wave paradigm was more similar to the active task (involved active movement using the same button box), although increased activity has also been reported in previous studies (Berlot et al., 2019; Wiestler et al., 2011). To further mitigate concerns regarding differences in activity, and to ensure that any subsequent analysis would not be impacted by the main effect of task activity, for each participant we normalised activity levels for each task/digit, based on the highest activity value obtained for that digit (across clusters). These normalised values were used for the subsequent analysis.

Figure 3B&C shows the resulting activity gradient. For both active and passive tasks, activity within each cluster decreases as a function of distance from the target digit (with some variations relating to D1 and D5 overlap, due to double D1 representations, see Figure 2A). To characterise somatotopic activity gradients, difference in activity levels was calculated by subtracting mean activity values for each of the 10 digit-pairs across the 5 digit-selective clusters (see Figures 4A and S4) and taking the absolute average value. Higher numbers therefore indicate greater differences in activity levels. We then clustered these selectivity difference scores into meaningful somatotopic relationships for further comparison. Digit pairs were classified based on two main factors: whether or not the pair contained a digit target to its cluster (target versus non-target), and whether or not the digit pair comprised of neighbouring versus non-neighbouring digits. As an example, within target cluster D3, the digit pair of D3-D2 is classified as target-neighbour (see Figure S4 for an illustration of classification). Note: for this analysis, D5 was considered as a non-neighbour to D1, due to the prominence of “double-thumb” representations (i.e., two maps of the thumb in the SI hand map: Kikkert et al., 2016), this analysis was repeated with D5 as a neighbour to D1 – see supplementary methods.

**Figure 4:**
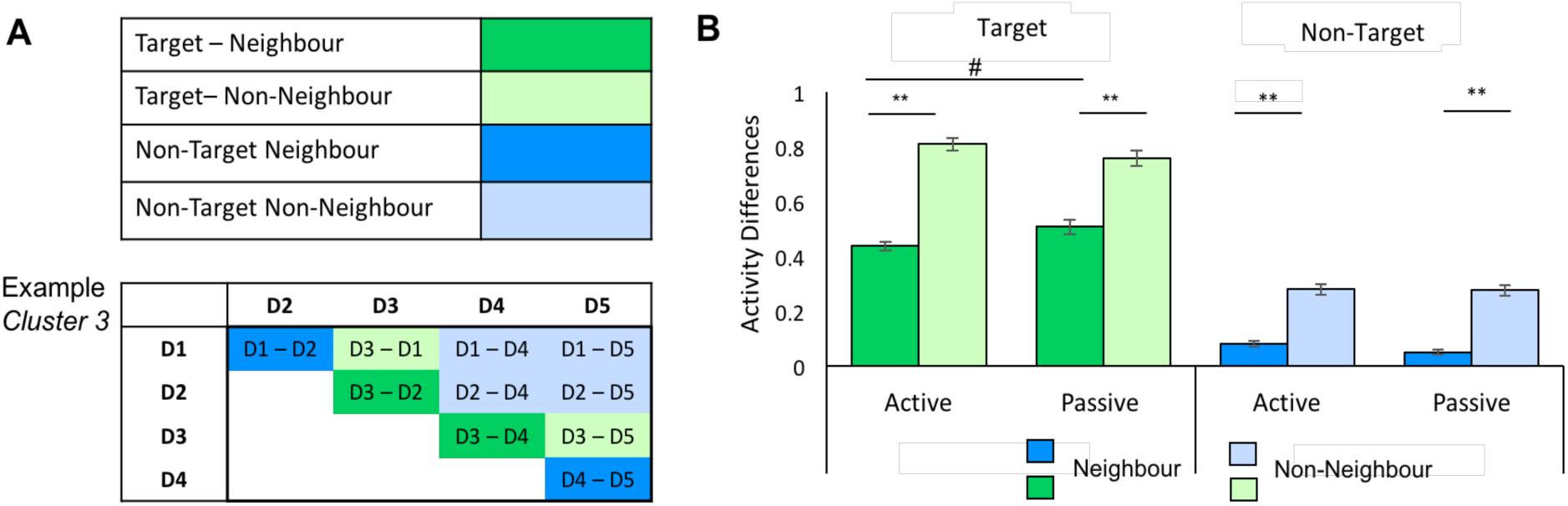
Comparing somatotopic profiles across the active and passive tasks. In this analysis, we characterised neighbourhood in the highly-selective digit-specific clusters. (A) Within each digit-specific cluster, pair-wise digit activity levels were contrasted. Each digit pair was characterised based on two criteria: whether it contains a neighbouring or a non-neighbouring digit pair (dark vs light colours); and whether the digit pair contained the target digit to that cluster (i.e. the digit with a corresponding affiliation to that of the cluster; green versus blue). Example for each of these four cases is shown for cluster 3 (see Figure S4 for all clusters). Results from this analysis are show for the active and passive tasks in (B). In all cases, we find that selectivity (shown here as activity differences) was greater for non-neighbouring digits, with respect to immediate neighbouring digits, which is a hallmark of somatotopic mapping. When considering somatotopic gradients relating to the target digit, we find that selectivity with respect to the immediate neighbours tended to be higher in the passive condition compared with the active condition, driving a significant interaction with respect to the tasks (and also with respect to tasks and nativity). ** indicates significant differences at p<.001, # indicates a trend.

To identify differences in somatotopic gradients across the tasks, a 2×2×2 repeated measure ANOVA was carried out with factors target (target vs. non-target digits), neighbourhood (neighbouring vs. non-neighbouring digits), and task (active vs. passive task). The 3-way interaction between these factors was significant: F(1,14)=43.116, p<.001, indicating that the somatotopic target/neighbourhood relationships differ across the two tasks. To interpret these differences, we followed-up with two 2×2 repeated measures ANOVAs to investigate the somatotopic gradient in the active and passive condition separately for the target and non-target digits.

For the target digits there was a significant 2-way interaction between neighbourhood and task: F(1,14)=32.798, p<.001. This appears to occur because, although the selectivity gradient between neighbouring and non-neighbouring digits is largely preserved across tasks, the selectivity profile is more pronounced in the passive task (figure 4B)– resulting in greater activity differences, specifically for the neighbouring digits. This was confirmed in the follow-up Wilcoxon matched-pair signed-rank tests: for both tasks, non-neighbouring digits have larger differences in activity, than neighbouring digits (Z = -*3.408*, p<.001; Figure 4B), therefore indicating a clear somatotopic gradient within each task. However, differences between target digits and their neighbours tended to be larger in the passive than in the active task (Z=-2.135, p=.033; note the adjusted alpha level for this test was .0125 to account for 4 comparisons). The difference between the target digit and non-neighbouring digits was not different between the two tasks Z=-.912, p=.362.

For completeness, we also looked at somatotopic relationships between digits that were not target to the digit selective cluster, for example within digit cluster 3, activity difference between neighbours (average D1-D2, D4-D5) was compared with non-neighbours (average D1-D4, D2-D5 etc.). For the non-target digits, there was no difference in somatotopic neighbourhood relations between tasks. This was indicated by the 2×2 repeated measures ANOVA, which did not produce a significant interaction F(1,14)=1.179, p=.296, nor a significant main effect of task F(1,14)=1.194 p=.293). Somatotopic relationships were, however, supported for non-target digits, as seen in a significant main effect of neighbour vs non-neighbour F(1,14)=250.501, p<.001, with non-neighbouring digits having larger differences in activity than neighbouring (Wilcoxon matched-pair signed rank Z=-3.412, p<.001; Figure 5B).

**Figure 5:**
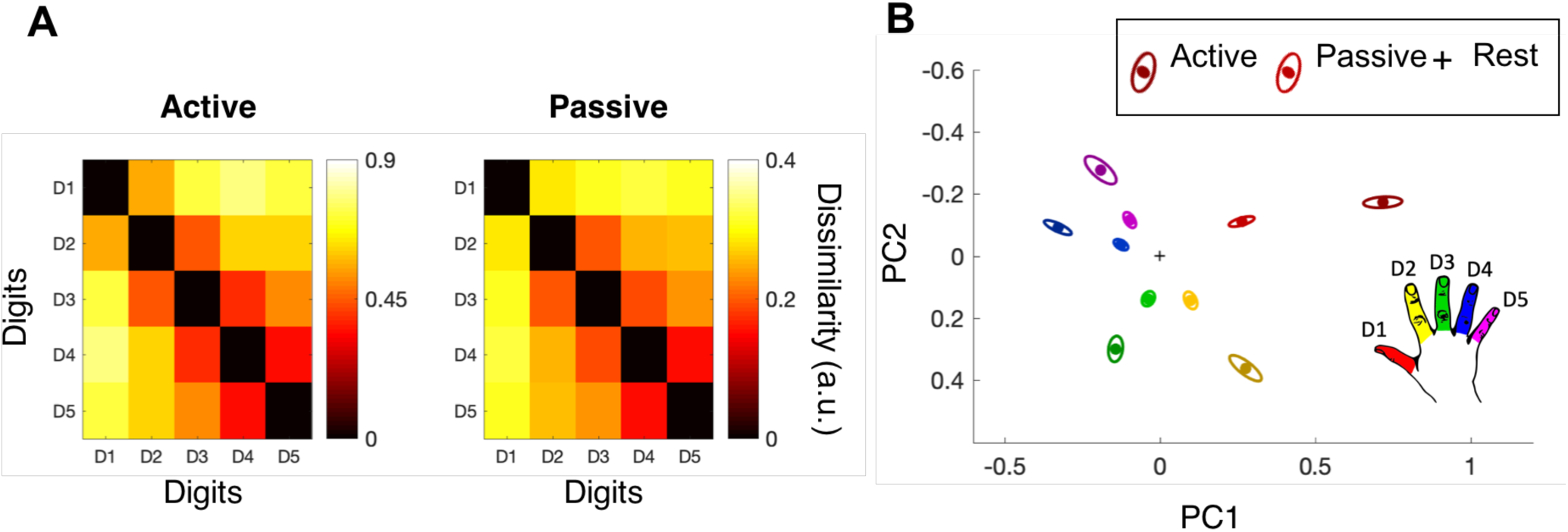
Multivariate hand representation is similar during active and passive tasks. (A) Group mean representational dissimilarity matrix for the active and passive tasks. The colour bars indicate dissimilarity such that hotter colours indicate greater dissimilarity across digit pairs. The dissimilarity scale was greater for the active task, indicating greater separability of patterns across digit pairs. (B) 2D depiction of the data in A, using multidimensional scaling. Ellipses indicate between-participant standard error, calculated separately per task. Red = D1, yellow = D2, green = D3, blue = D4 and pink = D5, darker colours represent the active task, whereas lighted colours represent the passive task. PCs: Principal components maximising inter-digit dissimilarity.

In sum, supporting our predictions, in both the active and passive condition there was a strong somatotopic gradient within each digit-specific cluster, with non-neighbouring digits showing larger differences to the target digit compared with neighbouring digits. We found that the passive task produced a slightly steeper somatotopic gradient with respect to the target digit and its neighbours (though not the target digit and its non-neighbours).

### 3.3 Multivariate pattern analysis confirms comparable hand representation across tasks

A complementary means to interrogate hand representation is the use of multivariate analysis such as representational similarity analysis (RSA; Ejaz et al., 2015; Nili et al., 2014)). For each pair of digits, the (unthresholded) activity patterns across the entire SI hand mask are compared, resulting in representational distances that are greater when the two digit patterns are more dissimilar. Therefore, the distance indicates digit selectivity. Unlike the dice analysis discussed above, this approach is independent of threshold. Indeed, RSA has been found to provide measures of representational similarity even in the absence of variation in activity levels (Dempsey-Jones, Wesselink, Friedman, & Makin, 2019; Diedrichsen, Wiestler, & Krakauer, 2013). It is also dissimilar to the ROI analysis discussed above in that this analysis is based on an anatomically-defined mask, freeing us from the above mentioned confound of task similarities between the functional localiser and the active task.

Figure 5A shows the inter-digit distance matrices calculated for the active and passive task. We first examined the mean distances across all digit-pairs (i.e. across each matrix) and compared them across tasks. Mean distance was significantly greater for the active than the passive task (t(13) = 10.9, p<.000). Consequently, as illustrated in Figure 5B, each digit is represented more independently of the others (illustrated with greater physical distances) in the active task. In other words, there is more information for inter-digit individuation, or digit selectivity, in the active task.

A further measure of functional hand representation is the inter-distance pattern across the 10 unique digit pairs, i.e. correlation between one representational dissimilarity matrix (RDM) and another. As shown in Figure 5A distances vary across digit pairs, resulting in relatively greater similarity between neighbouring to non-neighbouring digit pairs, reminiscent of the digit selectivity gradient shown in Figure 4B. The canonical hand representation has been shown to have strong consistency both at an intra- and inter-participant level (Ejaz et al., 2015). We, therefore, looked at how much individuals varied from this canonical pattern within either task. To do so, we calculated the correlation coefficients between each participant’s RDM and the average RDM obtained from all other participants for that task. This was done within-task, separately for active and passive conditions (e.g., participant_X_ active & average active), i.e. *intra*-task correlation. When comparing these two values (active mean rho = .90, passive mean rho = .91) we found no significant differences (t(13) = 0.16, p=.873), indicating that both tasks produce similarly high consistency across participants with respect to the group’s mean. Next, we tested for *inter*-task correlations, e.g., participant_X_ active & average passive, and vice versus from passive to active. If both tasks evoke a distinct representational structure, inter-task correlations should be lower than intra-task correlation. As summarised in Table 2, the correlations were not significantly different (assessed by paired t-tests), indicating the structure of the RDM is relatively stable independent of task demands.

**Table 2.** Both within and between task correlation values and associated T-Tests.

| Between-task correlations | Within-task correlations |  |
| --- | --- | --- |
|  | Active with active mean (mean rho = .90) | Passive with passive mean (mean rho = .91) |
| Active with passive mean (mean rho = .89) | $t(14) = 0.89$ ; $p = .388$ | $t(13) = 0.39$; $p = .706$ |
| Passive with active mean (mean rho = .90) | $t(13) = 0.21$; $p = .834$ | $t(13) = 0.69$; $p = .503$ |

## 4 Discussion

Here we compared digit activity created by two tasks: one involving passive cutaneous stimulation of the digit distal pad, and the other involving whole digit active movements. Despite marked differences between afferent input, efferent output and top-down factors, we show that the activity patterns that comprise the canonical somatotopic organisation is largely consistent in SI between these two tasks. This is demonstrated in the preservation of three key fundamental principles of somatotopy: spatial consistency of active and passive digit maps, a somatotopic gradient of activity levels for both tasks, and similar multivariate representational structure between tasks.

Notwithstanding the preservation of these critical features at both the macro-level (somatotopic map), and the meso-structure of fine-grained features (multivariate analysis), we did identify some differences between active and passive tasks. Specifically, consistent with previous reports (Berlot et al., 2019; Wiestler et al., 2011), univariate analysis demonstrated that the active task produced an overall greater amount of activity, compared to the passive task. We also found that, in the active task, the activity patterns generated by different digits were more clearly distinguishable. That is, on average there was greater separability of the multivariate activity patterns for the active than passive task (see below for more discussion). Somewhat inconsistently, our univariate activity analysis produced some evidence suggesting greater digit selectivity in the passive task. This selectivity advantage, however, was seen only for the comparison of target digits with their neighbours (and not with non-neighbours), and did not survive corrections for multiple comparisons – and will thus not be interpreted further. Overall, these between-task differences suggest active tasks may confer an advantage when disambiguation of the individual digits is required.

### 4.1 General consistency in spatial location of maps

Establishing the comparability between tasks is non-trivial, considering how extensively the primate hand map is studied – across fields, species and methods. Previous studies attempted to bridge these noticeable gaps by matching across task demands as closely as possible (Berlot et al., 2019). Here, by using tasks not tightly matched in stimulation features, we allow stimulation to vary between tasks in a manner more typical of, for example, variation between studies. Indeed, historically, differences in tactile stimulation protocols are often cited as the cause of divergent results, though largely without formal substantiation. For example, in the field of human brain remapping following amputation, traditional research using *passive* stimulation to the lower face has documented massive shifts of facial representation into the missing hand cortex (Flor et al., 1995). Other paradigms using *active* facial movements indicate relatively stable representation, with little activity in the missing hand area from facial movements (Kikkert, Johansen-Berg, Tracey, & Makin, 2018; Makin et al., 2013), and shifts of representation documented only locally within the face area (Makin et al., 2015; Raffin, Richard, Giraux, & Reilly, 2016). Some have interpreted the presence or absence of spatial position shifts for the facial map in the light of different stimulation protocols (Andoh, Milde, Tsao, & Flor, 2018; Flor, Diers, & Andoh, 2013), e.g., due to different underlying processes such as the engagement of different cortical layers in the SI microcircuit (Petreanu et al., 2012; though see Philip, Valyear, Cirstea, & Frey, 2017; Striem-Amit, Vannuscorps, & Caramazza, 2018; more on cortical layers below). However, our results here suggest that active and passive tasks elicit spatially overlapping somatotopies. Moreover, we showed comparable activity gradient for the non-target digits. This seems to suggest consistent activity gradients that are smaller than the spatial extent of the body-part itself. Still, we note that the spatial overlap between the active and passive maps was not complete (as indicated by the Dice values), indicating that there *is* some spatial displacement of maps between tasks. In sum, the relatively consistent spatial overlap of maps produced by active and passive tasks here seems to undermine arguments that differences in stimulation type are the sole cause of large scale differences in reported (re)organisation of the body map.

### 4.2 Active motor tasks and digit selectivity

Here, we found significantly more overall activity in response to active movement of a digit, as compared to passive stimulation of the tip of the digit. As mentioned in the Introduction, more activity could be the result of the greater number and range of afferent and top-down inputs associated with active movement. Even solely considering cutaneous afference, active tasks tend to produce more forceful indentations of the tips of the digits than light touch, air puffs or electrostimulation. The greater source of inputs in active tasks could contribute to greater signal-to-noise ratio and, in turn, more information to afford the increased inter-digit dissimilarity as found with the multivariate analysis. For example, Arbuckle and colleagues (2019) compared active digit movement at different speeds and showed that faster movement (i.e. more motor recruitment) induced greater inter-digit dissimilarity in the primary sensorimotor cortex.

It might also be that paradoxically the active motor task produces greater inter-digit separability than passive cutaneous stimulation because of inter-digit enslaving. This is because the information produced by an active task is more aligned with typical, ecologically relevant patterns of sensory input. It has long been known from electrophysiological work in primates that later somatosensory areas (such as BA 1 and 2) contain neurons with whole hand or multi-digit representations (reviewed in Iwamura, Tanaka, Iriki, Taoka, & Toda, 2002)). Earlier areas (and BA 3b in particular) – while containing predominantly single-digit receptive-field (RF) neurons – also contain a host of more complicated RF structures, including multiple points of excitation on the same digit, and multi-digit RFs (Iwamura, Iriki, & Tanaka, 1994; Thakur, Fitzgerald, & Hsiao, 2012). Considering the ubiquity of motor-related input to the SI hand map, Hebbian plasticity subsequent to motor activity may cause these complex RF properties (Overduin, d’Avella, Carmena, & Bizzi, 2014) see also (Dempsey-Jones et al., 2019; Dempsey-Jones et al., 2016; Ejaz et al., 2015 for related findings in humans). Since active movement involves more complex inputs from across the hand – this may produce a more optimal input to these multi-digit RFs. Considering that in daily life single digits are rarely tactually target in complete isolation, this account advocates for usage of ecologically-valid tasks to optimally activate SI.

A different consideration relates to the neuroimaging acquisition technique used here and other related studies, namely EPI. Research shows that typical (gradient-echo) EPI sequences are more sensitive to activity of the superficial layers (Goense, Merkle, & Logothetis, 2012). When considering the SI microcircuit, different cytoarchitectonic layers of the cortex contain neurons that preferentially respond to afferent inputs and efferent outputs, with the superficial layers (II/III) receiving efferent information that will be more relevant in the context of the active task (Friston, 2010; Thomson & Bannister, 2003; Yu et al., 2019). Relatedly, work from London and Miller (2013) has shown that neurons in BA 2 have different timing responses to inputs from active and passively induced movements. Therefore, while being largely co-localised within an area, active and passive tasks may produce inputs that are differently timed– but this cannot be differentiated due to the sluggish hemodynamic response relied upon by fMRI. Finally, here we examined digit representations across multiple sub-divisions of SI (namely areas 3b, 3a and 1). While some characteristics are shared across these sub-divisions and they are tightly linked (in particular areas 3b and 1; Burton & Fabri, 1995; Friedman, Chen, & Roe, 2004; Kaas, 1983), as mentioned above, there are also distinct characteristics within each of these areas. Therefore, it may be possible that active and passive tasks may produce distinct digit representations within specific sub-divisions. For example, in areas with predominantly single-digit RFs, passive tasks may be optimal. Indeed, Berlot and colleagues (2019), previously demonstrated a non-significant trend towards increased separability of digits using passive stimulation in areas 3b and 1.

### 4.3 Sensory gating and efference copies

An important reason why somatosensory processing may be different during active and passive tasks, is because sensory systems need to be able to distinguish whether sensory input is self-generated (through volitional movement) or externally-generated. Computational models of motor control stipulate that this is achieved by the motor system sending information to the sensory system about the expected sensory outcomes of a movement (efference copy; Wolpert & Flanagan, 2001). The influence of efference copies on somatosensory processing have been suggested to be either excitatory, through ‘corollary discharge’ mimicking the expected sensory feedback, or inhibitory, through cancellation or gating of somatosensory responses during movement (London & Miller, 2013).

Classical research on sensory gating has demonstrated that perception of tactile stimuli (Blakemore, Frith, & Wolpert, 1999; Shergill, Bays, Frith, & Wotpert, 2003) and processing of sensory input (Seki, Perlmutter, & Fetz, 2003) is suppressed during voluntary movements. Within somatosensory cortex, less activity in has been documented during active tasks (Cohen & Starr, 1987; Voss, Ingram, Haggard, & Wolpert, 2006; Wasaka, Hoshiyama, Nakata, Nishihira, & Kakigi, 2003), in particular when sensory feedback of a voluntary movement matches the actual feedback (Blakemore, Wolpert, & Frith, 1998). However, other research found that voluntary movements produce equal somatosensory activity, whether or not the movement lead to tactile stimulation (Blakemore et al., 1998; Jansma, Ramsey, & Kahn, 1998). Additionally, single unit recordings have shown increased neuronal firing in somatosensory cortex prior to movement onset, in line with the idea of corollary discharge (Krupa, Wiest, Shuler, Laubach, & Nicolelis, 2004; London & Miller, 2013; Soso & Fetz, 1980). Layer-specific neuronal recordings indicate these early responses do not arise from ascending thalamic input, but rather may reflect modulation from motor cortex (Krupa et al., 2004). London and Miller (2013) reconcile these diverging findings by demonstrating different neurons in monkey SI showing different response properties with respect to active versus passive movements, with some showing increased activity, and others showing decreased activity. This suggests that both sensory gating and corollary discharge occur within SI during active movements.

Here, we find that overall activity is increased during the active task, which is not consistent with the idea of sensory suppression. While in our study this may be confounded by our active functional localiser, previous studies have also reported increased activity during active tasks (Berlot et al., 2019; Wiestler et al., 2011). We also found significantly increased pattern separability for the active task compared with the passive. These differences found between active and passive tasks could be in part also driven by motor system influence, in the form of efference signals, on sensory processing.

### 4.4 Conclusions

Our study suggests active and passive tasks may be similarly used for exploring the somatosensory hand representation using fMRI. This has practical consideration for the implementation of studies on the ground – as active tasks are easier to set up, and don’t require as much specialist equipment. It has further practical implications for situations where either active or passive stimulation may not be possible: such as in people with limited mobility in the former case, or amputees who are missing physical digits but can produce active movements in their phantoms, in the latter case (Kikkert et al., 2016). On a more fundamental level, our findings have deeper implications for our understanding of SI, as the product of a tight, and bidirectional relationship with the motor system.

## Acknowledgements

TRM was funded by a Sir Henry Dale Fellowship jointly funded by the Wellcome Trust and the Royal Society (grant number 104128/Z/14/Z) and an ERC Starting Grant (grant number 0032-2-289-6121). The authors would like to thank Sanne Kikkert for her assistance in conceptualising the project and for her assistance with analysis.

