## Supplementary Materials for "Similar somatotopy for active and passive digit representation in primary somatosensory cortex"

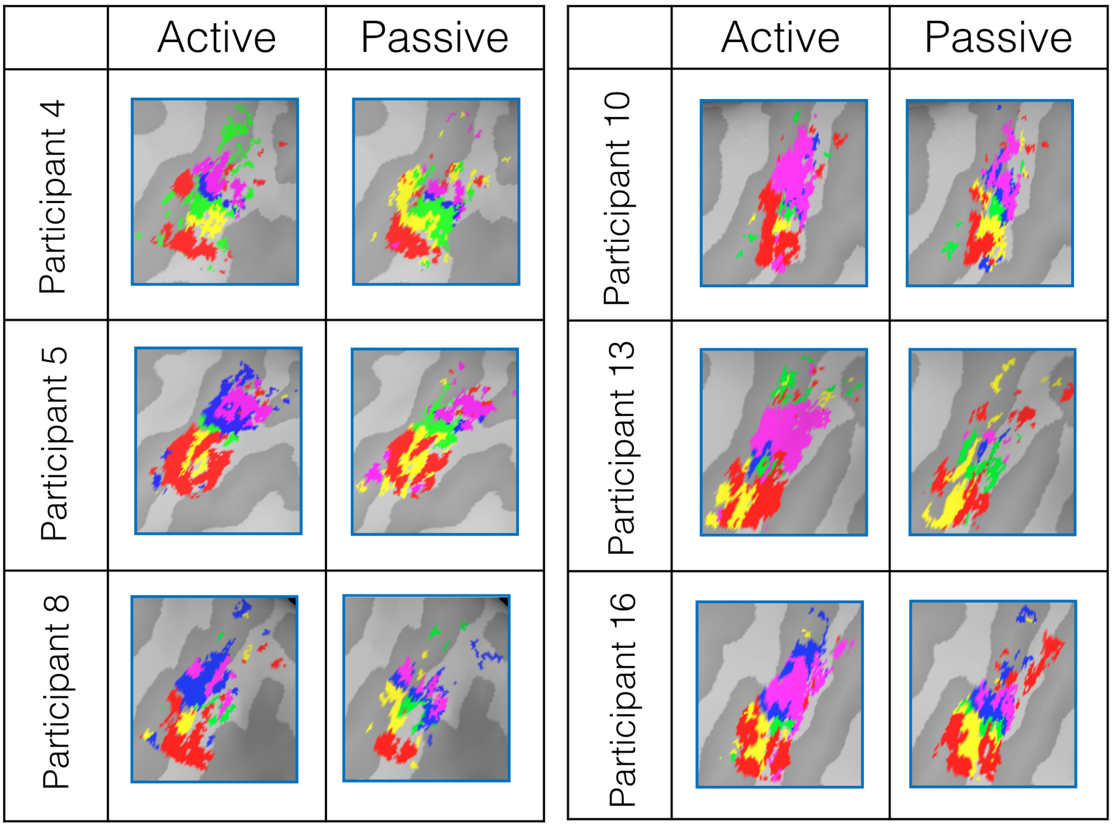

*Figure S1*: Minimally-thresholded activity within the SI hand mask of each digit for the active and passive task shown for six example participants. This data was used for the Dice analysis.

### Traditional Dice produces same results

The traditional Dice analysis is calculated by dividing the amount of spatial overlap between, in this case, two digit representations, by the total spatial area. Due to the variance in spatial extent across tasks and across digits, for the main analysis we divided the overlap by the smallest digit representation area, therefore normalising overlap to the smallest digit area. However, we also conducted the traditional Dice analysis and results were largely the same, despite lower overlap values on the whole (see figure S2). Dice values for the same digit across tasks decreased from .660 to .401, and for neighbouring digits from .310 to .176.

However, despite decreases in the average Dice coefficient values, the relationship between tasks and digit relations remained the same. Meaning that across the tasks, the same digit produces higher spatial overlap than each digit compared to its neighbour $t(14)=12.09, p<.001$ and compared to non-neighbouring digits $t(14)=11.23, p<.001$. Therefore, results were the same using the traditional dice analysis as using the modified dice analysis presented in the main text.

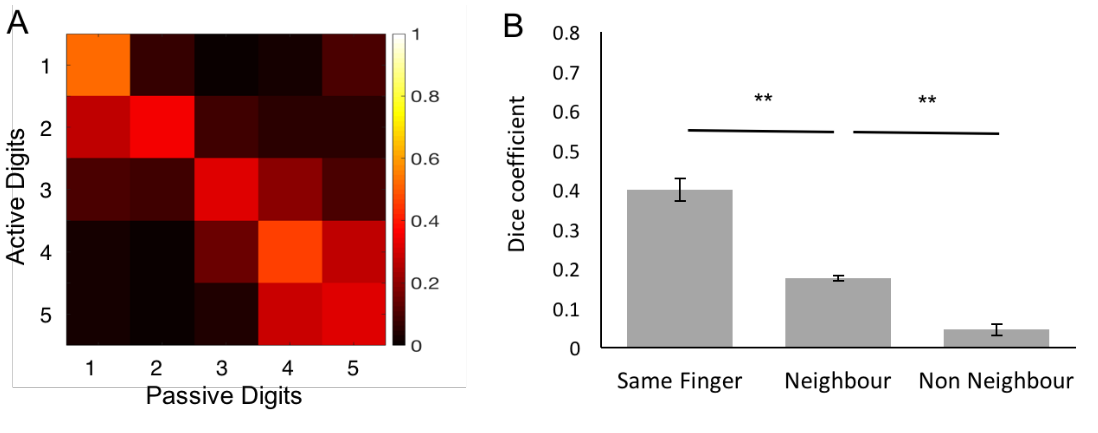

*Figure S2*: A) 5x5 matrix showing spatial overlap in SI hand mask as measured by the (non-normalised) dice coefficient between active and passive digits. The Dice coefficient shows greater overlap between homologous digits across tasks, as well as greater overlap between neighbouring than non-neighbouring digits. All other information is detailed for Figure 2.

**Somatotopic gradients are maintained when including D5 as a neighbour to D1**

When analysing the somatotopic gradient within digit-specific clusters, in the main text D5 was treated as a non-neighbour to D1. However, double D1 representation is a fairly common phenomenon (see figure 2 and SI). When conducting the same analysis as described in section 2.6.2 but including D5 as a neighbour to D1 (instead of as a non-neighbour), the main results remained largely unchanged. The 3-way ANOVA (2x2x2) showed a significant interaction $F(14,1)=22.628, p<.001$. The follow-up 2x2 ANOVA using just target digit differences also still showed a significant interaction $F(14,1)=16.547, p=.001$. Follow-up Wilcoxon related-samples signed-rank tests revealed that the differences between target and neighbouring digits was significantly smaller than differences between target and non-neighbouring digits for both active and passive tasks (both p=001). Additionally, the differences between target and neighbouring digits across tasks was also significantly different Z=-2.131, p=.033, with the passive condition having greater differences compared to active, although this finding does not survive multiple comparison correction. Finally, the difference between target and non-neighbouring digits was not significantly different between the two conditions Z=-.626, p=.531.

The 2x2 ANOVA just looking at non-target digit pairs did not show a significant interaction F(14,1)=.211, p=.653, nor a significant main effect of condition F(14,1)=.475, p=.502, but did show a significant main effect of neighbour vs non-neighbour F(14,1)=277.552, p<.001. Follow-up tests using Wilcoxon related-samples signed rank tests showed again that the difference between neighbouring digits was smaller than between non-neighbouring digits Z=-3.41, p=001.

These results were all in line with the results reported in the main analysis, indicating that including D5 as a neighbour to D1 does not affect the results of this analysis.

**Using a more conservative threshold for digit-specific masks produces same results**

In order to demonstrate that the somatotopic gradients also held at a more conservative FDR threshold, the analysis was repeated using digit-specific clusters based on an FDR threshold of .0001. Cluster sizes are decreased using this more conservative threshold as can be seen in supplementary table S1. However, when conducting the same analysis as described in section 2.6.2 the results remain largely unchanged.

| Digit | .0001 | | | .01 | | |
|---|---|---|---|---|---|---|
| | Average | Min | Max | Average | Min | Max |
| 1 | 415.2 | 92 | 771 | 606.7 | 200 | 1125 |
| 2 | 225.1 | 76 | 425 | 286 | 102 | 470 |
| 3 | 176.9 | 21 | 388 | 231.2 | 37 | 454 |
| 4 | 322.4 | 42 | 609 | 398.9 | 89 | 688 |
| 5 | 181.53 | 1 | 368 | 259 | 1 | 519 |

Table S1: Average, minimum and maximum number of voxels for each digit-specific cluster using FDR threshold of .0001 and .01.

The 3-way ANOVA (2x2x2) showed a significant interaction $F(1,14)=51.651$, $p<.001$. The follow-up 2x2 ANOVA using just target digit differences also still showed a significant interaction $F(1,14)=50.422$, $p<.001$. Follow-up Wilcoxon related-samples signed-rank tests revealed that the differences between target and neighbouring digits was significantly smaller than differences between target and non-neighbouring digits for both active and passive tasks ($Z = -3.408$, $p=.001$). The differences between target and neighbouring digits across tasks was also significantly different $Z=-2.385$, $p=.017$, with the passive condition having greater differences compared to active, although this does not survive correction for multiple comparisons (adjusted alpha for four comparisons = .0125). Finally, the difference between target and non-neighbouring digits was not significantly different between the two conditions $Z=-.454$, $p=.650$.

The 2x2 ANOVA just looking at non-target digit pairs did not show a significant interaction $F(1,14)=.1.007$, $p=.333$, nor a significant main effect of task $F(1,14)=.872$, $p=.366$, but did show a significant main effect of neighbour vs non-neighbour $F(14,1)=316.535$, $p<.001$. Follow-up tests using Wilcoxon related-samples signed rank tests showed again that the difference between target and neighbouring digits was smaller than between target and non-neighbouring digits $Z=-3.408$, $p=001$.

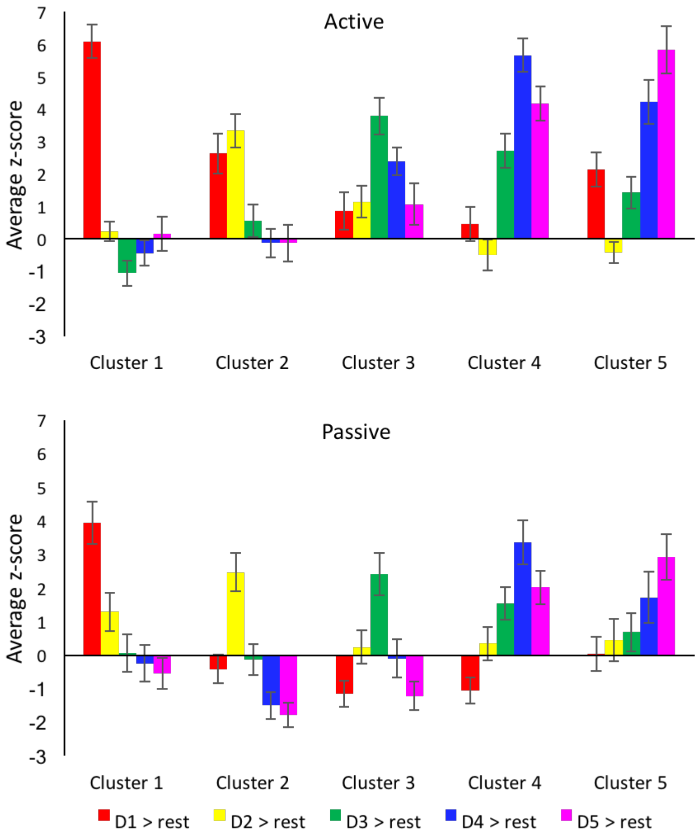

*Figure S3*: Average raw z-score values (before normalisation) for each digit in each digit-specific cluster. It can be seen that activity values are lower in the passive task than in the active task. All other information is detailed for Figure 3

| |
|---|
| Selected – Neighbour |
| Selected – Non-Neighbour |
| Non-Selected Neighbour |
| Non-selected Non-Neighbour |

Cluster 1

| | D2 | D3 | D4 | D5 |
|---|---|---|---|---|
| D1 | D1 – D2 | D1 – D3 | D1 – D4 | D1 – D5$^+$ |
| D2 | | D2 – D3 | D2 – D4 | D2 – D5 |
| D3 | | | D3 – D4 | D3 – D5 |
| D4 | | | | D4 – D5 |

Cluster 2

| | D2 | D3 | D4 | D5 |
|---|---|---|---|---|
| D1 | D2 – D1 | D1 – D3 | D1 – D4 | D1 – D5$^+$ |
| D2 | | D2 – D3 | D2 – D4 | D2 – D5 |
| D3 | | | D3 – D4 | D3 – D5 |
| D4 | | | | D4 – D5 |

Cluster 3

| | D2 | D3 | D4 | D5 |
|---|---|---|---|---|
| D1 | D1 – D2 | D3 – D1 | D1 – D4 | D1 – D5$^+$ |
| D2 | | D3 – D2 | D2 – D4 | D2 – D5 |
| D3 | | | D3 – D4 | D3 – D5 |
| D4 | | | | D4 – D5 |

Cluster 4

| | D2 | D3 | D4 | D5 |
|---|---|---|---|---|
| D1 | D1 – D2 | D1 – D3 | D4 – D1 | D1 – D5$^+$ |
| D2 | | D2 – D3 | D4 – D2 | D2 – D5 |
| D3 | | | D4 – D3 | D3 – D5 |
| D4 | | | | D4 – D5 |

Cluster 5

| | D2 | D3 | D4 | D5 |
|---|---|---|---|---|
| D1 | D1 – D2 | D1 – D3 | D1 – D4 | D5 – D1$^+$ |
| D2 | | D2 – D3 | D2 – D4 | D5 – D2 |
| D3 | | | D3 – D4 | D5 – D3 |
| D4 | | | | D5 – D4 |

*Figure S4*: Somatotopic profiles were compared across active and passive tasks in digit-specific clusters. As mentioned in figure 4, in this analysis pairwise digit activity levels were contrasted within each cluster. Digit pairs were characterised based on two criteria: whether it contains a neighbouring or a non-neighbouring digit pair (dark vs light colours); and whether the digit pair contained the target digit to that cluster (i.e. the digit with a corresponding affiliation to that of the cluster; green versus blue). This characterisation is shown here within each digit-specific cluster. + indicates cells which were altered for the analysis including D5 as a neighbour to D1.
